# Effect of prolonged dehydration stress on the vector competence of *Aedes aegypti* for Mayaro virus

**DOI:** 10.1101/2025.06.16.660000

**Authors:** Abdul Wahaab, Jaime Manzano Alvarez, Hieu Tran Nguyen Minh, Joshua B. Benoit, Jason L. Rasgon

## Abstract

*Aedes aegypti* is a competent vector for a variety of mosquito-borne viruses including Zika, chikungunya, Mayaro, yellow fever, and dengue, which cause debilitating diseases in animals and humans. It is highly invasive and is widely distributed across Asia, Europe, Oceania, and the Americas. Climatic factors such as relative humidity (RH) can have substantial effects on mosquito biological characteristics and the dynamics of pathogen spread. Low RH leads to dehydration in mosquitoes causing modifications in behavioral and physiological responses pertaining to pathogen spread, such as host-seeking behavior and blood-feeding patterns. Here, we evaluated the effects of prolonged dehydration stress on Mayaro virus infection and vector competence in *Ae. aegypti* mosquitoes. Our findings suggest that prolonged dehydration stress following Mayaro virus infection alters viral dissemination dynamics in mosquitoes, indicating that humidity may modulate intra-vector viral dissemination potentially impacting vector competence and pathogen spread. The previously observed effects of higher feeding and altered survival and our current observations on altered vector competence suggest that the impact of dehydration on viral transmission is expected to be complex and will be crucial to understanding the dynamic disease patterns of mosquito-borne viruses across diverse climatic conditions.

## 1 Introduction

Vector-borne diseases (VBD) are transmitted by hematophagous arthropods, and cause over 700,000 deaths and account for more than 17% of annual human illnesses caused by viruses, parasites, and bacteria [1–8]. Mayaro virus (MAYV) is an emerging and remerging alphavirus that was initially isolated from febrile workers in Mayaro County, Trinidad in 1954 [9]. MAYV is a single-stranded positive-sense RNA virus with a genome size of approx. 11.7 kb. It is classified into three genotypes: Genotype-D (spread across several South American countries), Genotype-L (confined to Brazil) [10], and the least prevalent Genotype-N (which emerged in Peru in 2010) [11]. MAYV fever presents in humans as a mild to severe self-limiting illness, which typically begins with an abrupt onset of fever, arthritis/arthralgia, and a maculopapular rash. Other symptoms include muscle pain, headache, nausea, vomiting, retro-orbital pain, and diarrhea. This acute debilitating illness, analogous to the Chikungunya virus (CHIKV; Togaviridae, Alphavirus) infection, often results in prolonged arthralgia in over 50% of affected individuals [12, 13].

The primary vector responsible for transmitting MAYV is the canopy-dwelling mosquito *Haemagogus janthinomys* (Dyar, 1921) [9]. Multiple vector competence studies have shown that *Aedes aegypti*, *Ae. albopictus, Anopheles freeborni*, *An. gambiae*, *An. quadrimaculatus*, *An. atroparvus,* and *An. stephensi* are competent vectors for MAYV under laboratory conditions, [14–19] demonstrating the virus’ significant capability to be permissive to multiple mosquito species. Over the years, MAYV has triggered localized epidemics and sporadic outbreaks, particularly in communities working and residing in close proximity to the Amazon forest. However, the number of outbreaks has increased in the last decade due to an increase in anthropogenic activities such as landscape fragmentation and the factors that are likely driven by climate change [20–22].

Mosquitoes are highly impacted by humidity, rainfall, and temperature, which impacts survival, density, feeding behavior, reproduction, geographical distribution, and consequently, their potential to transmit pathogens. However, the effects of climate change on VBDs can be complex and multifaceted [23, 24]. Relative humidity is considered one of the key determinants impacting the distribution and population density of *Ae. aegypti* [25–28]. Mosquitoes encounter varying levels of relative humidity throughout their life span ranging from 20-100 % with varying levels indoors and outdoors [29–31]. During extreme heatwaves, which result in high temperatures and low humidity levels, mosquitoes encounter dehydration stress due to extreme vapor pressure deficits [32], triggering physiological and behavioral responses to maintain osmotic balance and hemostasis. These may include modifying their cuticle composition, seeking alternative microclimates, and modifying spiracle morphology to minimize water loss and dehydration [33–37]. If these mechanisms prove inadequate, mosquitoes intensify blood-feeding and host-seeking activities to acquire water and stay rehydrated, while molecular responses including the expression of heat shock proteins and the production of antioxidants help in alleviating dehydration damage (7, 22, 28). Ultimately, fluctuations in relative humidity can significantly affect mosquito survival, vector competence, and pathogen transmission, particularly as climate change escalates the frequency of severe weather events (19-21).

The effect of temperature fluctuations on mosquito-virus interaction and pathogen spread has been widely studied and well-documented over the last decade, however, the impact of relative humidity changes on these dynamics remains insufficiently understood [38–40]. Our previous work examined the effect of short-term RH shock (low RH for < 24h) on mosquito vector competence for MAYV. Here, this study was designed to understand and compare the effects of prolonged, continuous dehydration stress on Mayaro virus infection vector competence in *Ae. aegypti* mosquitoes by assessing different components of vectorial capacity such as survival and changes in Mayaro virus infection, dissemination, and transmission. The findings will advance our understanding of complex disease patterns of mosquito-borne arboviruses and provide valuable insights into how varying critical climatic conditions influence disease transmission dynamics.

## 2 Material and Methods

### 2.1 Mosquito rearing

For general rearing, *Ae. aegypti* (Liverpool strain) mosquito eggs were obtained from NIAID/NIH Filariasis Research Reagent Resource Center for distribution by BEI Resources (NR-48921), NIAID, NIH. Mosquitoes were reared and maintained at the insectary located at Millenium Science Complex, Pennsylvania State University (University Park, USA) in 30 × 30 × 30 cm cages, under 27°C ± 1 °C, a 12:12 hours light:dark cycle, and 80% ± 5 relative humidity (RH). Larvae were fed intact and ground koi pellets (Tetra Pond koi Vibrance; Tetra, Melle, Germany). Adult mosquitoes were maintained with 10% sucrose solution *ad libitum*. Adult females were allowed to feed on anonymous human blood (BioIVT, NY, USA) for reproduction purposes following a previously described membrane feeder protocol [14].

### 2.2 Cells and Virus stock

Vero cells (derived from African green monkey kidney; CCL-81, ATCC, Manassas, VA, United States) were maintained in a complete growth medium containing DMEM (Dulbecco’s modified Eagles medium), supplemented with 10% FBS (Fetal bovine serum) and 1% SP (penicillin and streptomycin)] in a 37°C incubator with constant supply of 5% CO2 (all reagents were purchased from Gibco, Thermo Fisher Scientific, Waltham, MA, United States). Mayaro virus genotype L strain BeAr505411 (BEI Resources, Manassas, VA, United States) was originally obtained from *Ha. janthinomys* mosquitoes in Para, Brazil, in 1991. The virus was diluted in DMEM to MOI (multiplicity of infection) 0.01 and inoculated on Vero cells for one hour, after which cells were washed with DMEM and incubated with 15 mL of complete growth medium and incubated for 24 to 36 hours until the appearance of cytopathic effect (CPE). Subsequently, the infectious supernatant was aliquoted and stored at −80 °C. The viral titer of the frozen stock was measured using FFA (focus forming assays) before their use in experiments.

### 2.3 Humidity Treatment Setup

Two humidity treatments (80% RH and 35% RH) were prepared at a constant temperature of 27°C. To reach 35% RH, chambers were crafted with plastic transparent containers and cups holding super-saturated solution of MgCl_2_ [41]. Relative humidity was monitored before and during experiments using two digital hygrometers (ThermoPro, Ontario, Canada) for each chamber.

### 2.4 Experimental Design

Five to seven day old female mosquitoes were anesthetized on ice and sorted into 20 × 30 × 20 cardboard cages, which were placed overnight in regular insectary conditions (80% ± 5 RH and 27°C). Mosquitoes were then fed once for 60 minutes on infected human blood spiked with infectious Mayaro virus (1 × 10^7^ ffu/mL) at a ratio of 1:1. An aliquot of infectious blood was collected in a 1.5 ml Eppendorf tube, centrifuged, and stored at −80°C for further titration. Fully engorged mosquitoes anesthetized on ice were sorted and placed in multiple 10 × 10 × 10 cm cages. The cages were randomly divided into two groups and exposed to either normal humidity conditions (80% ± 5 RH and 27°C) or reduced humidity conditions (35% ± 5 RH and 27°C) until samples were collected at 7 days post-infection (dpi) and 14 dpi, respectively. Mosquitoes in both groups were maintained with 10% sucrose solution ad libitum provided through a parafilm membrane that was changed every 24 hours. Mortality was recorded every day, and around thirty surviving mosquitoes were collected per humidity treatment group at each time point (7 and 14 dpi) for vector competence assays. Mosquito bodies (including wings and head), legs, and saliva were individually collected in mosquito-diluent and spitting solution, with each sample processed separately for each mosquito, as described previously [42]. The samples (body and legs) were homogenized by single zinc-plated steel, 4.5 mm bead (daisy, Rogers, ARE, USA) using a TissueLyser II (QIAGEN GmbH, Hilden, Germany) on 30 Hz for two two-minute cycles. Later, they were clarified by centrifugation and stored at −80°C for further experiments. The experiment was run in three independent experimental replicates and results of each replicated are reported individually. The layout of the experimental design is illustrated in Figure 1.

**Figure 1:**
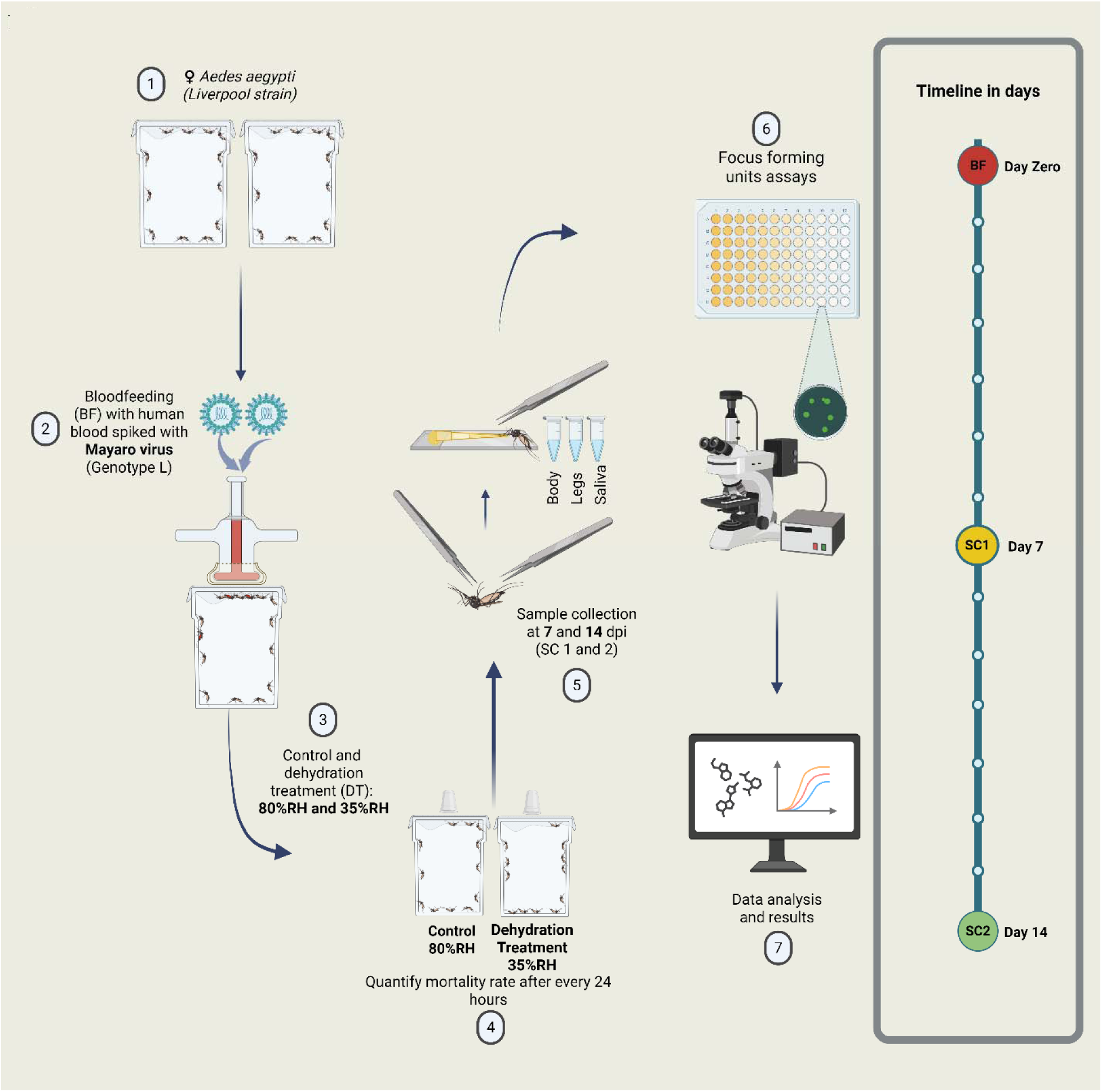
Graphical layout of the experiment design: The schematic demonstrates the experimental setup and overall workflow. The adult Aedes aegypti (Liverpool strain) female mosquitoes were fed with human blood spiked with the Mayaro virus (genotype-L). Fully engorged Mosquitoes were sorted and cohort into two different humidity conditions i.e 80% (standard insectary conditions) and 35% (dehydration treatment/relative humidity shock) with access to 10% sugar solution all the time. The mortality was observed every 24 hours and samples (Bodies, Legs, and Saliva) were collected at 7 and 14 days post-infection (dpi)respectively. The viral titers were assessed using Focus forming assays (FFA) and data was analyzed using GraphPad prism. The image was created in created in https://Biorender.com.

### 2.5 Vector Competence assays

Assayed mosquitoes were anesthetized with trimethylamine (Sigma Aldrich, St Louis, MO, United States) at their respective collection time points (7dpi and 14dpi). Legs were detached from the body, and mosquitoes were forced to salivate into a pipette tip with 1:1 mixture of FBS (fetal bovine serum) and 50% sugar solution. Body, legs, and saliva were collected in 2ml safe lock Eppendorf tubes (Hamburg, Germany) with 300, 300, and 100 µL of mosquito diluent respectively. The samples (body and legs) were homogenized (as described above) and stored at −80°C for viral titration. All the samples (body, legs, and saliva) were used to prepare a ten-fold dilution (10^2^ to 10^5^ for the body, 10^1^ to 10^4^ for the legs, and 10^0^ or 10^0^ to 10^1^ for saliva) for focus-forming assays (FFAs). Vector competence rates are reported as infection rates (IR) which stand as a proportion of infected bodies over the total, dissemination rates (DR) which stands for the proportion of infected legs over the infected bodies, transmission rates which stand as a proportion of infected saliva over infected legs and transmission efficacy (TE), which stands as the proportion of infected saliva over total collected mosquitoes.

### 2.6 Focus Forming assay (FFA)

The detection of live Mayaro virus (MAYV) infectious particles in samples from mosquito legs, bodies, and saliva was performed using FFAs in Vero cells. Cells were plated in 96 well (flat bottom) plates at a concentration of 4 × 10^4^ cells/well. After 24 hours, a series of ten-fold dilutions of the samples were prepared in serum-free DMEM, and 30 µL of the mixture was used to infect the cells at 37°C with 5% CO2 for 60 minutes. Subsequently, the supernatant was discarded and replaced by 100 µL of overlay medium (1:1 mixture of complete DMEM medium and 1.6% methylcellulose) and incubated for 24 hours at 37°C with 5% CO2. After 24 hours the cells were fixed with 4% paraformaldehyde for fifteen minutes, and washed with 1x PBS three times. Then the cells were permeabilized with 0.2% Triton X in 1XPBS for fifteen minutes and washed thoroughly again with 1XPBS. Viral particles were detected using the monoclonal anti-chikungunya virus E2 envelop glycoprotein clone CHK-48 (BEI Resources, Manassas, VA, United States) diluted 1:500 in 1XPBS solution as described previously [43]. The wells were washed three times with 1XPBS and the samples were incubated with the secondary antibody Alexa 488 goat anti-mouse IgG (Invitrogen, Eugene, Orlando, United States) at 1:2500 dilution with 1XPBS, followed by the final wash with 1XPBS (three times). Olympus BX41 inverted microscope equipped with UPlanFI 4× objective and FITC filter was used for counting and recording Mayaro virus foci.

### 2.7 Statistical analysis

Statistical analysis was performed using Graphpad Prism 10 for Windows, version 10.4.2 (633) (Graphpad Software, La Jolla, CA, United States). Viral titers in mosquito bodies, legs, and saliva were analyzed using the unpaired (non-parametric) t-test. Infection, dissemination, transmission, and mortality rates are illustrated as percentages and were analyzed using Fisher’s exact test. A “p-value” less than 0.05 (p < 0.05) was considered significant.

## 3 Results

### 3.1 Dehydration stress does not affect Mayaro virus infection in *Aedes aegypti* mosquitoes

Humidity levels notably impact mosquito behavior and the transmission of mosquito-borne pathogens [44]. To understand whether prolonged RH stress interferes with Mayaro virus infection dynamics, bodies (including head, abdomen, and wings) of the mosquitoes from both humidity treatment groups (i.e. normal kept at 80% humidity levels and RH stress which were kept at 35% humidity levels) were collected at 7 dpi (days post-infection) and 14 dpi in mosquito diluent, and viral titer was determined using FFA (focus-forming assay). Infection rates in the RH shock mosquitoes group were found to be similar to that of mosquitoes that were kept in normal humidity conditions at 7 and 14 dpi, with no statistically significant difference detected except in one replicate at 14 dpi (Figure 2A to 2F). Moreover, bodies also showed no statistically significant difference in viral titers at 7 dpi (Figure 2G; *p =* 0.811, 2H; *p =* 0.219, and 2I; *p =* 0.068) and 14 dpi (Figure 2J; *p =* 0.552, 2K; *p =* 0.997, and 2L; *p =* 0.563.) between both humidity treatment groups across all three replicates. These results indicate that transient environmental humidity fluctuations appear to have no impact on Mayaro virus infection dynamics in *Ae. aegypti* mosquitoes and were similar to our previous findings, in which short-term humidity shock had no significant effect on Mayaro virus infection [45].

**Figure 2:**
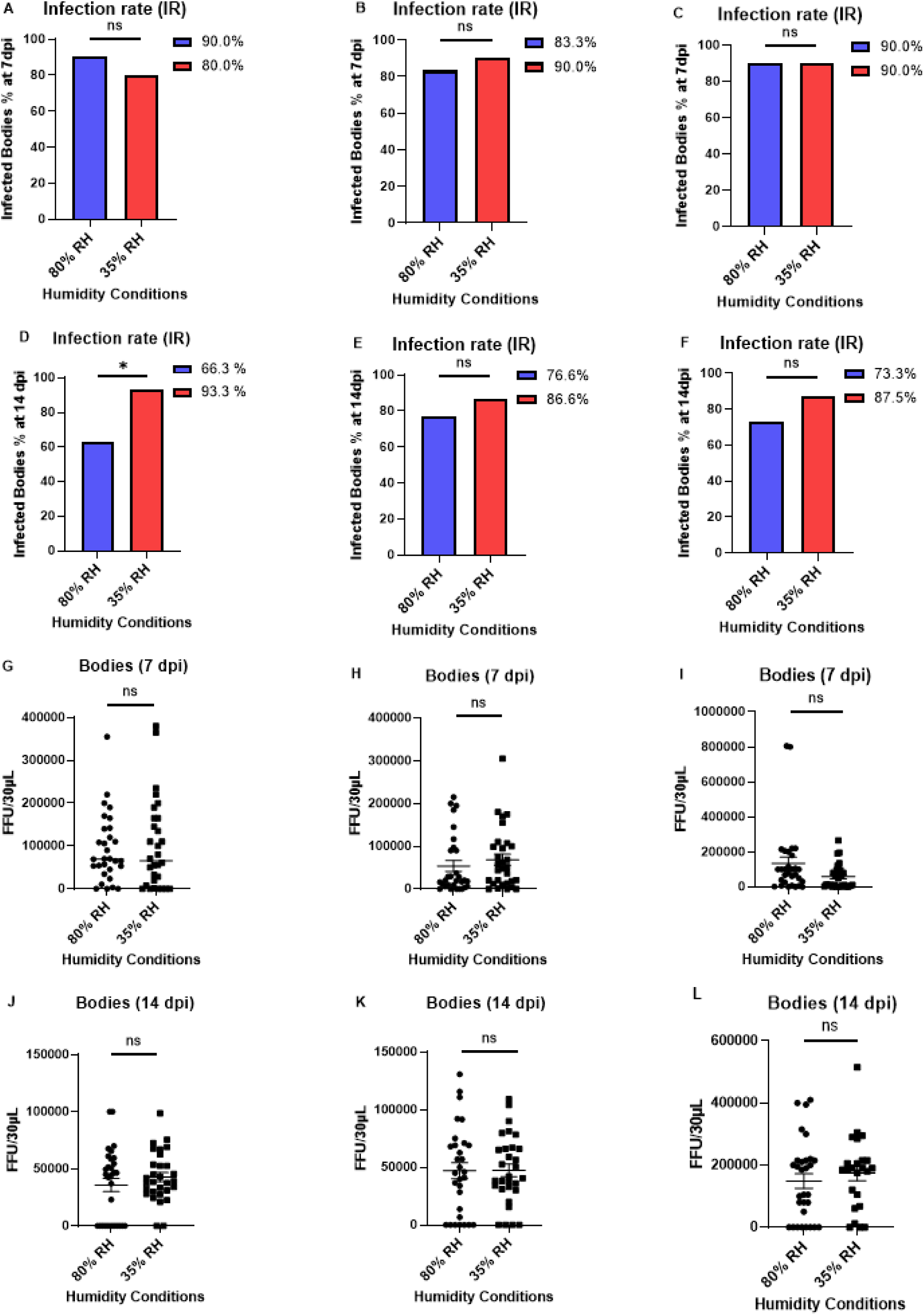
Infection rates and mosquitoes bodies viral titers: Infection rates (A to F) and bodies viral titers (G to L) were measured and compared in Mayaro virus-infected mosquitoes exposed to two different humidity conditions (i.e 80% and 35%) at 7 and 14 days post-infection (dpi). **A to F)** Infection rates are illustrated as percentages and were analyzed using a Fisher’s exact test at 7 dpi and 14 dpi. Subfigures A, B, and C show individual experimental replicates for infection rates at 7dpi, while D, E, and F show replicates at 14dpi. **G to L)** Viral titers in mosquitoes bodies are represented as focus forming units per 30 µL in (FFU/30 µL) and were analyzed using a non-parametric t-test at 7 dpi and 14 dpi. Subfigures G, H, and I show individual experimental replicates for viral titers in bodies at 7 dpi while J, K, and L show replicates at 14 dpi. Data illustrates three independent experimental replicates. Error bars demonstrate the standard error of the mean (SEM). Statistical difference is indicated where applicable. Statistical differences are presented as follows: ns (non-significant) for p > 0.05, and * for P < 0.05.

### 3.2 Dehydration stress modulates Mayaro virus dissemination in *Aedes aegypti* mosquitoes

To understand whether relative humidity (RH) shock interferes with Mayaro virus dissemination dynamics, the legs of the Mayaro virus-infected mosquitoes from both humidity treatment groups (80% & 35% RH) were collected at 7 dpi (days post-infection) and 14 dpi in mosquito diluent, and viral titer was determined using FFA (focus-forming assay). Dissemination rates in the RH stress group were found to be similar to that of mosquitoes that were kept in normal humidity conditions at 7 and 14 dpi, with no statistically significant difference detected among them in all three replicates (Figure 3A to 3F). However, a statistically significant increase in viral titers in the legs was detected at 7 dpi in one replicate (Figure 3I; *p =* 0.0176), and at 14 dpi in two independent replicates (Figure 3K; *p =* 0.0108 and Figure 3L; *p =* 0.0176), suggesting a potential pattern of higher viral dissemination in mosquitoes maintained under normal humidity conditions compared to those subject to dehydration stress or RH shock. These results indicate that transient environmental humidity fluctuations appear to impact Mayaro virus dissemination dynamics and are consistent with the studies that suggest that higher humidity levels may commonly associate with enhanced transmission potential of arboviruses by prolonging vector longevity [46–48].

**Figure 3:**
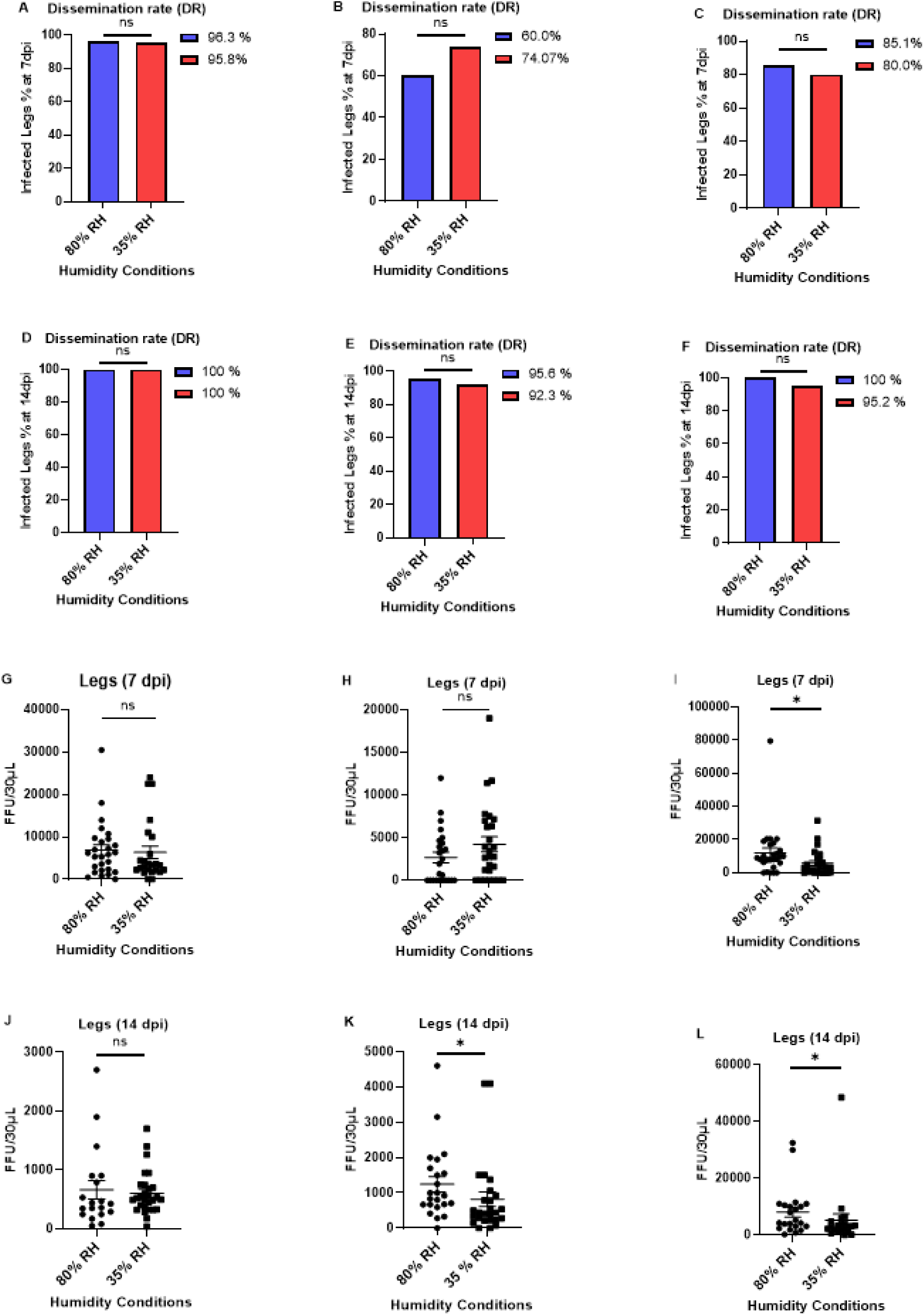
Dissemination rates and mosquitoes legs viral titers: Dissemination rates (A to F) and legs viral titers (G to L) were measured and compared in Mayaro virus-infected mosquitoes exposed to two different humidity conditions (i.e 80% and 35%) at 7 and 14 days post-infection (dpi). **A to F)** Dissemination rates are illustrated as percentages and were analyzed using a Fisher’s exact test at 7 dpi and 14 dpi. Subfigures A, B, and C show individual experimental replicates for dissemination rates at 7dpi, while D, E, and F show replicates at 14dpi. **G to L)** Viral titers in mosquitoes legs are represented as focus forming units per 30 µL in (FFU/30 µL) and were analyzed using a non-parametric t-test at 7 dpi and 14 dpi. Subfigures G, H, and I show individual experimental replicates for viral titers in legs at 7 dpi while J, K, and L show replicates at 14 dpi. Data illustrates three independent experimental replicates. Error bars demonstrate the standard error of the mean (SEM). Statistical difference is indicated where applicable. Statistical differences are presented as follows: ns (non-significant) for p > 0.05, and * for P < 0.05.

### 3.3 Dehydration stress does not affect Mayaro virus transmission in *Aedes aegypti* mosquitoes

To gain insight into whether relative humidity (RH) shock interferes with Mayaro virus transmission dynamics, the saliva of the Mayaro virus-infected mosquitoes from both humidity treatment groups (80% & 35% RH) were collected at 7 dpi (days post-infection) and 14 dpi in spitting solution, and viral titer was determined using FFA (focus-forming assay). Viral transmission rates (Figures 4A to 4F) and saliva viral titers (Figures 4G to 4L) in the RH stress mosquitoes were similar to that of mosquitoes that were kept in normal humidity conditions at both 7 and 14 dpi, with no statistically significant difference detected among them in all three replicates (Figure 4G; *p =* 0.1198, 4H; *p =* 0.342, 4I; *p =* 0.5697, 4J; *p =* 0.3305, 4K; *p =* 0.7601, and 4L; *p =* 0.8778). These results indicate that transient environmental humidity fluctuations appear to have a statistically indistinguishable impact on Mayaro virus transmission dynamics in Aedes aegypti mosquitoes. These results were similar to previous findings, where short-term humidity shock and lower humidity levels had no significant effects on Mayaro and Zika virus transmission dynamics, respectively [45, 49].

**Figure 4:**
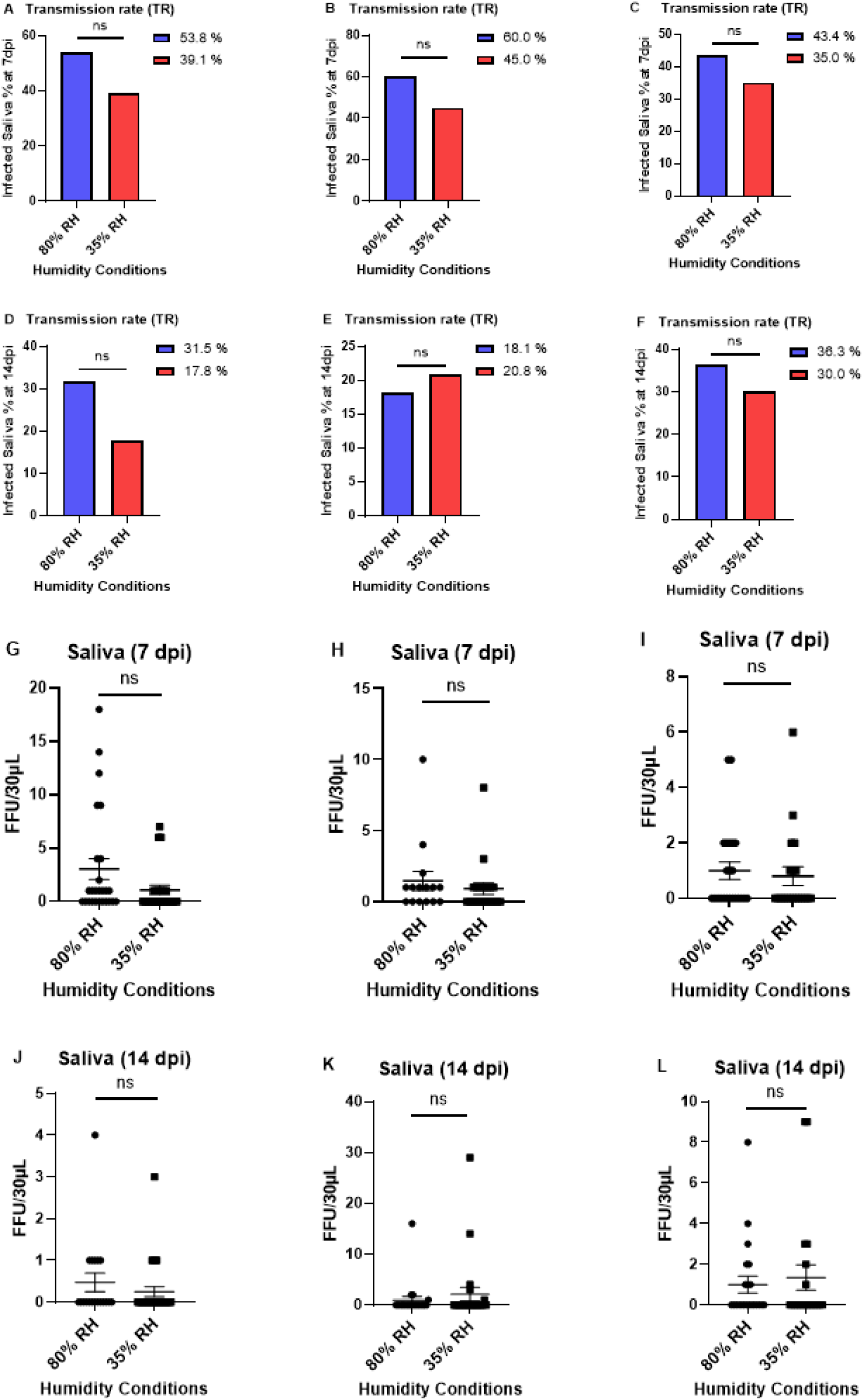
Transmission rates and mosquito saliva viral titers: Transmission rates (A to F) and saliva viral titers (G to L) were measured and compared in mosquitoes exposed to two different humidity conditions (i.e 80% and 35%) at 7 and 14 days post-infection (dpi). **A to F)** Transmission rates are illustrated as percentages and were analyzed using a Fisher’s exact test at 7 dpi and 14 dpi. Subfigures A, B, and C show individual experimental replicates for transmission rates at 7dpi, while D, E, and F show replicates at 14dpi. **G to L)** Viral titers in mosquit saliva are represented as focus forming units per 30 µL in (FFU/30 µL) and were analyzed using a non-parametric t-test at 7 dpi and 14 dpi. Subfigures G, H, and I show individual experimental replicates for viral titers in saliva at dpi while J, K, and L show replicates at 14 dpi. Data illustrates three independent experimental replicates. Error bars demonstrate the standard error of the mean (SEM). Statistical difference is indicated where applicable. Statistical differences are presented as follows: ns (non-significant) for p > 0.05, and * for P < 0.05.

### 3.4 Dehydration stress does not impact mortality rates following Mayaro virus infection in *Aedes aegypti* mosquitoes

Climate and weather may directly impact vector biology, including survival, longevity, biting, fecundity, and pathogen replication and development in the vector, which is critical for pathogen spread [50]. To investigate whether long-term relative humidity (RH) stress affects the mortality rates of Mayaro virus-infected mosquitoes, the mortality of the infected mosquitoes was observed independently at 7dpi and 14dpi using distinct cohorts of mosquitoes kept in separate boxes for each timepoint and humidity conditions (mortality rates are summarized in Table No. 1). The mortality rates in the RH stress group were found to be similar to that of mosquitoes that were kept in normal humidity conditions at both 7 and 14 dpi, with no statistically significant difference detected among them in replicates 1 and 2. However, a difference was seen in replicate 3, where a sharp increase in mosquito mortality was observed on 11 dpi (where 57.02% of total mosquitoes died in a single day) in the RH shock mosquitoes group, significantly raising their mortality rates from mosquitos kept in normal humidity conditions (which showed a gradual increase in mortality rates over time). This mortality spike, limited to the specific group and replicate, may reflect experimental variability, inconsistency in humidity chambers, and other unknown/uncontrolled factors, therefore, merits further exploration. These results collectively indicate that transient environmental humidity fluctuations appear to have no distinguishable impact on mortality rates following Mayaro virus infection in *Ae. aegypti* mosquitoes.

## 4 Discussion

The World Health Organization (WHO) considers health issues related to climate change among the greatest challenges of the 21st century. These changes, directly and indirectly, interfere with the natural environment, and when connecting them to climate and tropical diseases, it is evident that these fluctuations disrupt the ecosystem balance, contributing to an increase in the transmission of vector-borne diseases, with flaviviruses being among the most significant [51–53]. While the effects of temperature variations on mosquito-virus interactions and pathogen transmission have been well-documented over the past decade, the influence of relative humidity changes on these dynamics remains insufficiently understood. Preliminary entomological investigations on the effects of humidity on vectors suggest the significance of this variable [54–56]. In this study, we aimed to understand the physiological and virological consequences of prolonged dehydration stress on MAYV vector competence in *Ae. aegypti* by comparing mosquitoes maintained under normal humidity conditions (80% RH) with those exposed to prolonged dehydrating conditions (35% RH) post-MAYV infection. Key components of vectorial capacity, such as mortality, infection, dissemination, and transmission, were assessed to determine the impact of dehydration on viral transmission dynamics.

The non-infected *Ae. aegypti* mosquitoes, which were exposed to prolonged dehydration stress and had access to 10% sucrose solution provided on a cotton ball (enclosed in parafilm) all the time, did not show any significant mortally until 14 days (data not shown). A previous study assessed short-term dehydration effects following MAYV infection in mosquitoes and found higher mortality rates compared to the control group, suggesting that dehydration stress may increase the mosquitoes vulnerability during viral infection [45]. This has also been reported previously with other blood-feeding arthropod-borne viruses, and the stressors induced by environmental variables like thermal stress [57–59], possibly attributable to the need of resources for sustaining cellular hemostasis, virus-mediated modulation in gene expression, and the cost of immune responses activated in response to viral infection [60–62]. Primarily, dehydration can be severely stressful and requires discrete factors to maintain cellular hemostasis and facilitate recovery, which is likely disrupted during acute viral infection [32, 63, 64]. Here, we observed that the mosquitoes show no significant difference in mortality rates (Table No. 4) when they have access to sugar water, suggesting that long-term dehydration protocols did not impose a lethal physiological burden on mosquitoes, while also supporting the theory that blood feeding may help mosquitoes to prevent dehydration and its associated consequences despite the presence of viral infection [65]. The possible variation in mortality rates in other studies may also result from differences in dehydration exposure times and protocols, differences in blood sources used for mosquito feeding, mosquito species used, and other uncontrolled variables such as insectary and environmental conditions, or handling procedures.

We evaluated long-term impacts of dehydration on viral vector competence and the short-term impact of viral infection. Notably, we did not observe any significant variation in infection rates (Fig. 2a through f), dissemination rates (Fig. 3a through f), and transmission rates (Fig. 4a through f) between both groups at 7 and 14 days post-infection. Additionally, the viral titers in bodies (Fig. 2g through L) and saliva (Figure 4g through L) also showed no differences across both conditions, however, lower humidity or prolonged dehydration stress was associated with lower viral dissemination as evidenced by lower viral titers in the legs (Fig. 3g through L). This suggests that environmental stress (like lower humidity levels) may restrict the mosquito’s ability to support or transport the virus beyond the midgut (initial infection site) to other secondary tissues, potentially limiting the mosquito’s capacity to transmit the virus to the host. This impairment may also result from dehydration-induced physiological changes impacting midgut barrier integrity or viral replication efficacy and escape. Despite reduced dissemination, the consistent infection rates in the midgut and virus in saliva suggest that while dehydration may have compromised internal viral spread, it does not hinder initial viral establishment in the midgut or the infectious release of particles in saliva. It is plausible that a subset of mosquitoes still sustains sufficient viral replication to support transmission despite physiological stress [66–68].

Multiple studies have reported that viral vector competence can be adversely affected under scenarios that trigger stress in mosquitoes, including fluctuations in dynamic climatic conditions, particularly temperature [38–40, 69]. Since dehydration triggers physiological modifications in mosquitoes [32, 64, 70–72], we speculated that long-term dehydration could also modulate the vector competence and viral transmission dynamics, which may not exactly align with the outcomes of this study. However, intermittent but recurring exposure to low-humidity conditions has been reported to directly impact mosquito reproduction and physiology [71], suggesting that future studies may want to target how chronic and repeated bouts of dehydration may impact or deteriorate the mosquito’s immune system and increase vulnerability to viral infection, thereby modulating the viral transmission dynamics. Another direction of future research could be to investigate how dehydration stress and limited water availability influence mosquito feeding patterns, and consequently, viral transmission potential. Previous studies have shown that dehydration increases host-seeking behavior and blood-feeding frequencies [63, 73], specifically during the periods of low water availability and nutrient depletion [71]. Importantly, secondary feeding has been observed due to dehydration [73], and these additional blood meals have been shown to increase viral transmission [74].

Recent comparative studies of *Ae. aegypti* populations have demonstrated differences in blood feeding patterns and arbovirus transmission linked with climatic conditions and urbanization levels [75, 76], reflecting that dehydration and virus transmission dynamics may vary across *Ae. aegypti* lineages. MAYV has also been documented to infect multiple *Anopheles* mosquito species both *in vitro* and *in vivo* [77–79] and some earlier studies have elucidated the effects of relative humidity on *Anopheles* mosquito biology [80, 81]. Future investigations should also designed to examine how genetic variation in both arboviruses and mosquito species or sub-lineages interface with humidity and other climatic factors to influence viral transmission dynamics.

## 5 Conclusion

Our finding demonstrates that prolonged exposure to dehydration stress after Mayaro virus infection results in altered viral dissemination dynamics within *Ae. aegypti* mosquitoes. This suggests that humidity may modulate intra-vector viral spread, potentially impacting vector competence and pathogen spread. Beyond their immediate biological implications, these results also underscore the importance of recognizing the role of environmental conditions, which are crucial for refining predictive models for arbovirus spread, particularly in the context of a rapidly evolving climate.

## Acknowledgements

This work was funded by NIH/NIAID grant R01AI148551 to J.B.B. and J.L.R., and NIH/NIAID grant R01AI150251, USDA Hatch funds (4769), and funds from the Dorothy Foehr Huck and J. Lloyd Huck endowment to J.L.R.

## Supplementary Tables

**Table 1:** Summary of mortality rates: Mortality rates are measured and analyzed using a Fisher’s exact test at both 7 and 14 days post-infection (dpi). Data illustrates three independent experimental replicates. Statistical difference is indicated where applicable and are presented as follows: ns (Non-Significant) for p > 0.05, and * for P < 0.05.

| Mortality rates |  |  |  |  |  |  |  |  |  |  |
| --- | --- | --- | --- | --- | --- | --- | --- | --- | --- | --- |
| Replicates | Time point | 80 % Relative Humidity |  |  |  | 35 % Relative Humidity |  |  |  | Statistical Difference (P-Value) |
|  |  | Total Mosquitoes | Dead | Alive | Mortality Rates (%) | Total Mosquitoes | Dead | Alive | Mortality Rates (%) |  |
| Replicate 1 | 7dpi | 63 | 7 | 56 | 11.1 | 65 | 7 | 58 | 10.7 | Non-Significant (P > 0.9999) |
|  | 14 dpi | 63 | 33 | 30 | 52.3 | 65 | 24 | 41 | 36.9 | Non-Significant (P = 0.1091) |
| Replicate 2 | 7dpi | 65 | 04 | 61 | 6.1 | 57 | 02 | 55 | 3.5 | Non-Significant (P = 0.6838) |
|  | 14 dpi | 65 | 07 | 58 | 10.7 | 65 | 12 | 53 | 18.4 | Non-Significant (P = 0.3208) |
| Replicate 3 | 7dpi | 104 | 06 | 98 | 5.7 | 120 | 17 | 103 | 14.1 | *(Significant) (P = 0.042) |
|  | 14 dpi | 104 | 16 | 88 | 15.3 | 121 | 97 | 24 | 80.1 | ****(Significant) (P < 0.0001) |

**Supplementary Table 1:**
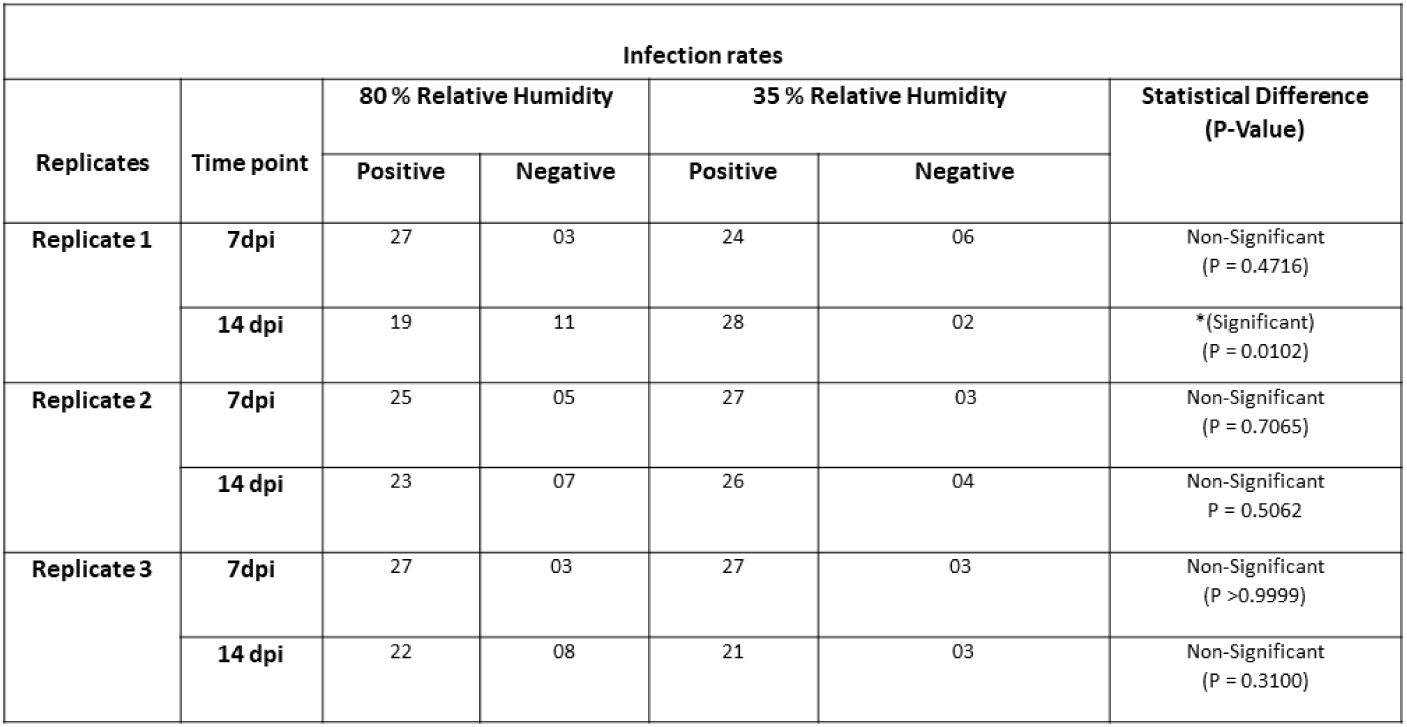
Summary of infection rates: Infection rates are measured and analyzed using a Fisher’s exact test at both 7 and 14 days post-infection (dpi). Data illustrates three independent experimental replicates. Statistical difference is indicated where applicable and are presented as follows: ns (Non-Significant) for p > 0.05, and * for P < 0.05.

| Infection rates |  |  |  |  |  |  |
| --- | --- | --- | --- | --- | --- | --- |
| Replicates | Time point | 80 % Relative Humidity |  | 35 % Relative Humidity |  | Statistical Difference<br>(P-Value) |
|  |  | Positive | Negative | Positive | Negative |  |
| Replicate 1 | 7dpi | 27 | 03 | 24 | 06 | Non-Significant<br>(P = 0.4716) |
|  | 14 dpi | 19 | 11 | 28 | 02 | *(Significant)<br>(P = 0.0102) |
| Replicate 2 | 7dpi | 25 | 05 | 27 | 03 | Non-Significant<br>(P = 0.7065) |
|  | 14 dpi | 23 | 07 | 26 | 04 | Non-Significant<br>P = 0.5062 |
| Replicate 3 | 7dpi | 27 | 03 | 27 | 03 | Non-Significant<br>(P >0.9999) |
|  | 14 dpi | 22 | 08 | 21 | 03 | Non-Significant<br>(P = 0.3100) |

**Supplementary Table 2:** Summary of dissemination rates: Dissemination rates are measured and analyzed using a Fisher’s exact test at both 7 and 14 days post-infection (dpi). Data illustrates three independent experimental replicates. Statistical difference is indicated where applicable and are presented as follows: ns (Non-Significant) for p > 0.05, and * for P < 0.05.

| Dissemination rates |  |  |  |  |  |  |
| --- | --- | --- | --- | --- | --- | --- |
| Replicates | Time point | 80 % Relative Humidity |  | 35 % Relative Humidity |  | Statistical Difference<br>(P-Value) |
|  |  | Positive | Negative | Positive | Negative |  |
| Replicate 1 | 7dpi | 26 | 01 | 23 | 01 | Non-Significant<br>(P >0.9999) |
|  | 14 dpi | 19 | 0 | 28 | 0 | Non-Significant<br>(P >0.9999) |
| Replicate 2 | 7dpi | 15 | 10 | 20 | 07 | Non-Significant<br>(P = 0.3775) |
|  | 14 dpi | 22 | 01 | 24 | 02 | Non-Significant<br>(P >0.9999) |
| Replicate 3 | 7dpi | 23 | 04 | 20 | 05 | Non-Significant<br>(P = 0.7224) |
|  | 14 dpi | 22 | 0 | 20 | 01 | Non-Significant<br>(P = 0.4884) |

**Supplementary Table 3:** Summary of transmission rates: Transmission rates are measured and analyzed using a Fisher’s exact test at both 7 and 14 days post-infection (dpi). Data illustrates three independent experimental replicates. Statistical difference is indicated where applicable and are presented as follows: ns (Non-Significant) for p > 0.05, and * for P < 0.05.

| Transmission rates |  |  |  |  |  |  |
| --- | --- | --- | --- | --- | --- | --- |
| Replicates | Time point | 80 % Relative Humidity |  | 35 % Relative Humidity |  | Statistical Difference<br>(P-Value) |
|  |  | Positive | Negative | Positive | Negative |  |
| Replicate 1 | 7dpi | 14 | 12 | 09 | 14 | Non-Significant<br>(P = 0.3932) |
|  | 14 dpi | 06 | 13 | 05 | 23 | Non-Significant<br>(P = 0.3121) |
| Replicate 2 | 7dpi | 09 | 06 | 09 | 11 | Non-Significant<br>(P = 0.4998) |
|  | 14 dpi | 04 | 18 | 05 | 19 | Non-Significant<br>(P >0.9999) |
| Replicate 3 | 7dpi | 10 | 13 | 07 | 13 | Non-Significant<br>(P = 0.7556) |
|  | 14 dpi | 08 | 14 | 06 | 14 | Non-Significant<br>(P = 0.7499) |

## Notes

### Competing Interest Statement

The authors have declared no competing interest.

